# Fine-scale spatiotemporal predator-prey interactions in an Antarctic fur seal colony

**DOI:** 10.64898/2026.04.03.716266

**Authors:** Ane Liv Berthelsen, Johannes Bartl, Alexander Winterl, Cameron Fox-Clarke, Jaume Forcada, Rebecca Nagel, Ben Fabry, Joseph Ivan Hoffman

**Author notes:** Corresponding author: Ane Liv Berthelsen. Department of Evolutionary Population Genetics, Faculty of Biology, Bielefeld University, Konsequenz 45, 33501 Bielefeld, Germany. Joint first authors (Ane Liv Berthelsen, Johannes Bartl). Joint senior authors (Ben Fabry, Joseph Ivan Hoffman).

## Abstract

Density is a major determinant of population dynamics, with high densities exacerbating intraspecific competition and disease transmission, while low densities increase predation risk. To investigate spatiotemporal density patterns and predator-prey interactions in a breeding colony of Antarctic fur seals (*Arctocephalus gazella*), we deployed an autonomous camera capturing minute-by-minute, high-resolution images throughout a breeding season. Using a YOLO-based neural network, we identified adult males, females and pups, as well as three avian predator-scavengers: giant petrels (*Macronectes* spp.), brown skuas (*Stercorarius antarcticus*) and snowy sheathbills (*Chionis alba*). Analysis of 4.1 million automated detections from over 10,000 high-quality images revealed spatiotemporal abundance patterns corresponding with the known breeding and foraging behaviours of these species. Strong temporal associations emerged between the abundance of pups and two avian species, while fine-scale spatial analyses showed that pups grouped together with other pups and adult females but avoided avian predators and territorial males. Notably, proximity to adult fur seals of both sexes reduced pup predation risk, defined as the distance between the pup and the nearest bird, whereas proximity to other pups did not. Overall, this study provides a framework for quantifying density-dependent interactions in wild populations and emphasises the value of remote observation in ecological research.

## 1. Background

Density strongly influences individual fitness and population persistence (1), either positively or negatively. Negative density dependence constrains population growth through intensified competition for resources, increased disease transmission and heightened predation (2–4). Conversely, positive density dependence, or an Allee effect, enhances population growth via the benefits that individuals gain from group living (5). These include increased mating success, cooperative feeding, improved predator detection due to shared vigilance (6) and enhanced predator deterrence due to the intimidating effect of large groups (7). Additionally, the dilution effect (8) lowers individual predation risk in large groups, as predators can only attack a limited number of prey. However, these mechanisms only operate above a critical density threshold, below which Allee effects no longer occur and survival declines (9).

Empirical evidence for an Allee effect was recently found in Antarctic fur seals (*Arctocephalus gazella*) at Bird Island, South Georgia (10). Here, two neighbouring breeding colonies–the special study beach (SSB) and freshwater beach (FWB)–differ in social density by a factor of four, yet breeding females from both localities most likely forage in the same areas and have access to the same food resources (11,12). Historically, population regulation in both colonies was mainly driven by negative density dependence, with higher pup mortality at high densities caused by traumatic injuries from territorial males and starvation following mother-pup separation or abandonment (11,13). However, a recent demographic decline attributed to climate-driven reductions in the availability of the seals’ staple diet, Antarctic krill (*Euphausia superba*), has steadily reduced the number of animals coming ashore (14). As a result, pups born at the low density colony (FWB) now experience higher mortality, largely due to predation, compared with pups born at the high-density colony (SSB) (10).

On land, Antarctic fur seals have few predators, with the pups being the main targets of predation (15). Documented instances of active predation involve giant petrels (*Macronectes: M. halli* and *M. giganteus*) and brown skuas (*Stercorarius antarcticus*), both of which are opportunistic predator-scavengers (16,17). Recent observations uncovered two distinct predation techniques used by giant petrels: pecking and drowning (15). During attacks, they strike from the ground and often begin feeding on the pup while it is still alive (15). In addition to preying on live pups, both species rely heavily on fur seal carrion and placentae during their breeding seasons (16,17). Snowy sheathbills (*Chionis alba)* are also frequently present in fur seals colonies (18). Although they typically associate with penguin colonies, where they kleptoparasitise by stealing regurgitated krill (19), sheathbills are flexible foragers that readily scavenge fur seal placentae and carcasses (20), but only rarely prey on pups (18). Thus, these three avian species represent a gradient from active predation to scavenging, with giant petrels and brown skuas opportunistically switching between scavenging and active predation on pups, while snowy sheathbills act almost exclusively as scavengers (21,22).

The Antarctic fur seal breeding season begins in early November, when the adult males arrive to establish territories (23). During this period, predators have limited foraging opportunities. About a month later, the adult females arrive to give birth to their pups, which are conceived the previous season (24). On average, females give birth two days after coming ashore and mate around a week later, after which they alternate foraging trips at sea with nursing their pups ashore (24,25). With the arrival of the adult females and pups, the breeding colonies become rich feeding grounds for scavengers and predators, with weak and unattended pups being especially vulnerable to predation (10,18).

Breeding in high-density areas can in principle reduce predation risk, as predicted by Hamilton’s selfish herd theory, which posits that individuals in less dense areas face the greatest danger (26). In addition, Antarctic fur seal mothers have been observed defending their pups against predators (10), whereas adult males tend to ignore the pups, which sometimes leads to accidental trampling (18). Consequently, a pup’s predation risk depends not only on its own physical condition but also on its position relative to other conspecifics, including adults of both sexes and other pups (15,27).

As part of the long-term monitoring programme of the British Antarctic Survey (BAS), a designated study colony of Antarctic fur seals at Bird Island has been continuously monitored since 1981. Individual-and population-level data including visual counts of permanently identified individuals have enabled reconstruction of this population’s demographic trajectory over the past 90 years, providing insights into the effects of changing food availability on vital rates and demographic trends (14,28). However, while these data are invaluable, they provide limited resolution of fine-scale population density patterns and their influence on predator-prey interactions. This gap could be addressed through time-lapse imaging, a non-invasive and labour-efficient alternative to field-based observations. The still imaging of pinnipeds has previously been used to monitor abundance (29,30), estimate body mass (31) and analyse social distancing (32). However, to our knowledge, time-lapse imaging has not yet been applied to study predator-prey interactions in pinnipeds at fine spatial or temporal scales.

In this study, we deployed an autonomous time-lapse camera at the low-density colony, FWB. Over 56 days, the camera recorded 66,645 images at 1-minute intervals. A custom-trained neural network was used to identify and locate animals within a quality-filtered subset of the images, allowing us to quantify the temporal and spatial abundance of Antarctic fur seals and their associated avian predator-scavengers. We hypothesised that if these birds rely on Antarctic fur seal pups for sustenance, (i) bird and pup abundance would be positively associated over time and (ii) that their spatial distributions would strongly overlap. Using pup proximity to predators as a proxy for predation risk, we further examined fine-scale spatial associations between Antarctic fur seal pups, adult fur seals and the avian species. We predicted that (iii) pups would associate with adult females but avoid adult males and avian predators, in accordance with fur seal social behaviour and predator avoidance. Finally, to assess whether group-living reduces predation risk, we evaluated whether the proximity of conspecifics influences pup predation risk. Specifically, we hypothesised that if adult fur seals deter avian predators, (iv) proximity to adults–particularly adult females–would decrease predation risk by increasing the distance between pups and their nearest avian predators.

## 2. Methods

### (a) Study site

The study was conducted at FWB, an Antarctic fur seal breeding colony at Bird Island, South Georgia (54°00024.800 S, 38°03004.100 W) (Figure 1a, b) during the austral summer of 2020–2021. The colony lies directly in front of the BAS field base, allowing the convenient installation and maintenance of the remote observatory (see below). FWB is intersected by a shallow stream, which creates natural corridors and affects the spatial distribution of the seals within the colony.

**Figure 1:**
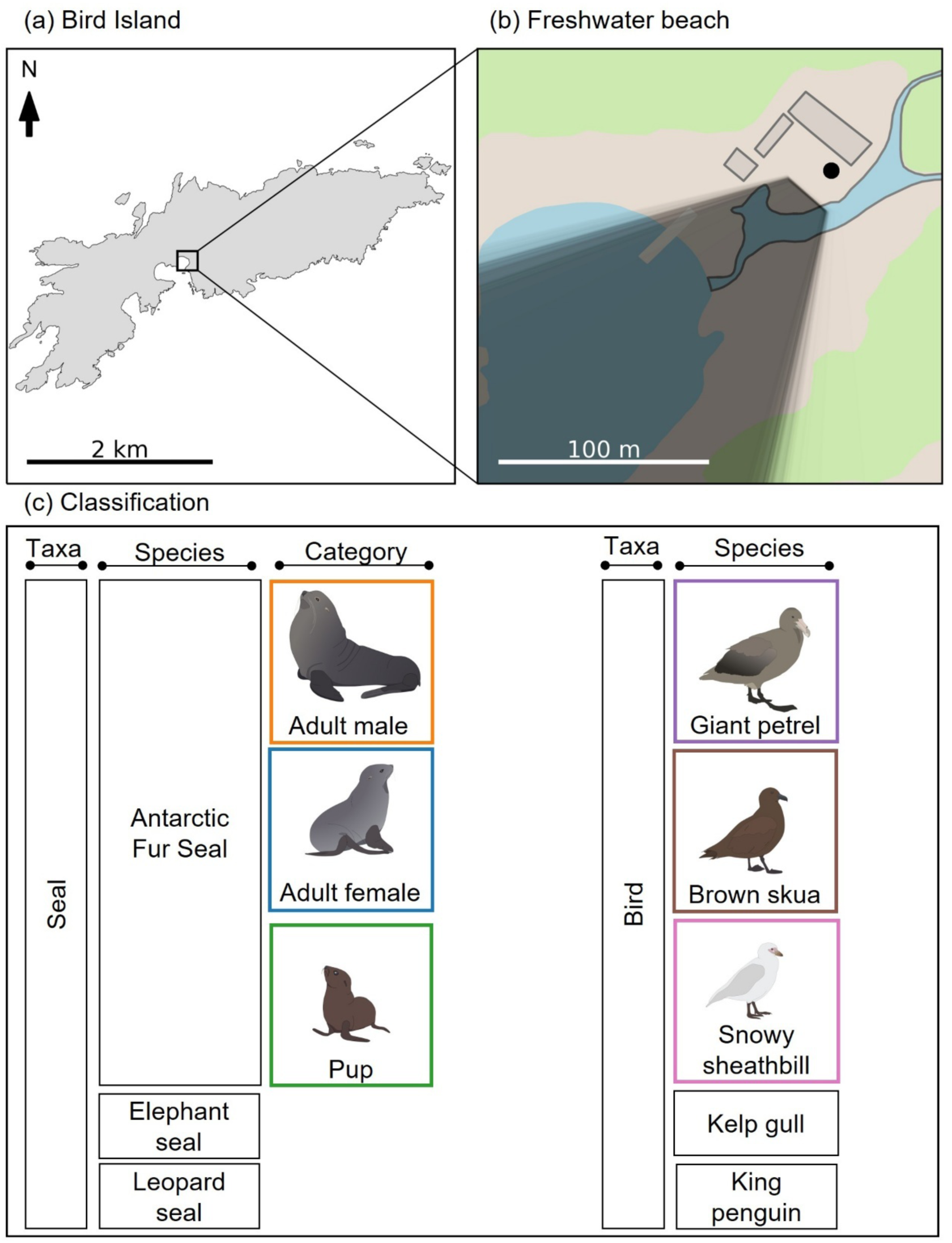
(a) Map of Bird Island, South Georgia; (b) The study colony (Freshwater beach, FWB) showing the fixed position of the camera (black point), which was mounted on the field station tower. The monitoring field-of-view (FOV) of the camera varied slightly over the course of the study period due to strong winds and manual exchange of SD cards. The grey area shows the FOV of the camera, where darker shading indicates more frequent observations compared to lighter shaded areas. (c) The classification hierarchy used for training the neural network, depicting the 13 classes (boxes) and highlighting the six focal classes (coloured boxes) of the study, with original artwork by ALB.

### (b) Remote observatory

We recorded images of the colony using an automated camera system (33) composed of a 16 mega-pixel digital mirrorless consumer-grade camera (Panasonic Lumix DMC G5) with a 14mm lens (horizontal field of view 23°), a controller unit based on an Arduino microcontroller, a tripod, a water-resistant housing, and a 12V car battery for the power supply. We mounted the camera on the field station’s tower at a height of eight meters and continuously recorded for 56 days between the 30th of October and the 24th of December 2020, spanning the peak of the Antarctic fur seal breeding season, at a rate of one image per minute, creating a dataset of 66,645 images.

### (c) Training dataset

We excluded recording days with poor visibility due to fog or precipitation (three days) from the dataset and selected at least one image for each of the remaining days (total = 53 images) for annotation. We selected these images to represent natural variation in the recording conditions, including differences in ambient light levels (from high to low), visibility, shadows and seal abundance. Using the image annotation tool Clickpoints (34), we manually annotated all observations of animals. Each observation was assigned a class – taxa, species or further differentiating category (for further information, see below) – and a rectangular bounding box enclosing the whole animal. To accurately reflect biological reality in dense aggregations where individuals frequently rest against one another, our annotation explicitly allowed for overlapping bounding boxes. In cases of image occlusion, boxes were placed to encompass the individual’s full estimated outline as if it were fully visible (see SI Figure 1). These bounding box annotations recorded the class, position in the image and size information, which were then used to train a neural network for the automated detection of different classes by size and appearance.

In these 53 training images, we observed individuals of the following species: Antarctic fur seal, elephant seal (*Mirounga leonina*), leopard seal (*Hydrurga leptonyx*), king penguin (*Aptenodytes patagonicus*), kelp gull (*Larus dominicanus*), giant petrel, brown skua and snowy sheathbill. Where species classification was not possible, we labelled bird and pinniped species as ‘bird’ and ‘seal’, respectively. We further subdivided the Antarctic fur seals into adult males, adult females and pups, unless differentiation was not possible (0.4 % of observations). This yielded a total of 13 classes (bird, seal, eight species and three categories of Antarctic fur seal, Figure 1c). A total of 23,195 manually annotated observations were made.

### (d) Automated detection

For the analysis of the full dataset, we trained a neural network based on a EfficientNet B7 backbone (35) and a YOLO object detection head (36), implemented in python using the modules TensorFlow (37) and Keras (38), which supplied the initial weights (internal neural network parameters) for the network. The YOLO object detection head splits the image into a grid of boxes, called anchor boxes, where one box can contain one animal. As the grid spacing defines the granularity of detection, multiple such grids can be overlaid to handle different orders of object sizes. Based on the distribution of object sizes in the dataset of manual observations, we choose grid spacings of 64, 32 and 16 pixels. In each anchor box, the network predicts objects as a vector containing entries for its probability of being an animal, its position, width, height, and class, a structure resembling our bounding box annotation. We then split the manually annotated dataset consisting of 53 images (4608×3456 px) into a test dataset (12 images, 22.6 %) and training dataset (41 images, 77.4 %). To process the high-resolution source images, we utilized a data generator during training. This generator dynamically extracted 128×128-pixel crops from the training images and applied small random augmentations (e.g., random horizontal flipping, rotation, and scaling) to increase data diversity. We optimized the weights of the neural network with the optimization algorithm Adam (39) in two steps: a transfer learning step, where we optimized only the YOLO head weights and kept the EfficientNet backbone, and a fine-tuning step, where we optimized all weights.

The loss function used for optimization was identical to the published YOLO loss (36) except for the term handling the classes. The class error term in the original loss is the L2-distance (Euclidian distance) between a true and a predicted vector with one entry per class, which is one if the object belongs to the class and otherwise zero (one-hot encoding). The true vector will always contain only one non-zero entry, while the predicted vector contains a probability between one and zero for each class. This method cannot handle hierarchies in classes correctly. For example, an object manually labelled as *seal* but predicted by a neural network as a pup would result in a high loss. However, since the label is more general than the predicted subclass, we avoided penalizing such cases as follows: we adjusted the loss function by encoding the vector with a one for every subclass an object belongs to (e.g. *seal*, Antarctic fur seal, pup) and masking the L2-distance so that predictions that are more specific than the label (e.g. label: *seal* versus prediction: Antarctic fur seal) were not considered.

We trained the neural network on 128×128-pixel crops from the training set images for 10,000 training cycles during both the transfer learning and fine-tuning phases, with each cycle representing a full pass through the entire dataset, termed an epoch. Performance was evaluated on a set of 128 such crops after every epoch, and only the weight set with the minimal validation loss was kept for further analysis. We optimized the threshold for the accepted object probability of an anchor box to maximize the F1 score, a metric that balances prediction accuracy with how comprehensively relevant objects are detected. In addition, we applied a non-maxima suppression to suppress predicted bounding boxes overlapping by more than 75% and retained the one with the highest probability. Across the entire training dataset, we obtained an F1 score of 0.75. To assess performance variability, we also calculated F1 scores for each individual category (see SI Figures 2 and 3). These results confirmed the model’s capacity to make reliable predictions. We then applied the network to the whole dataset of image recordings.

**Figure 2:**
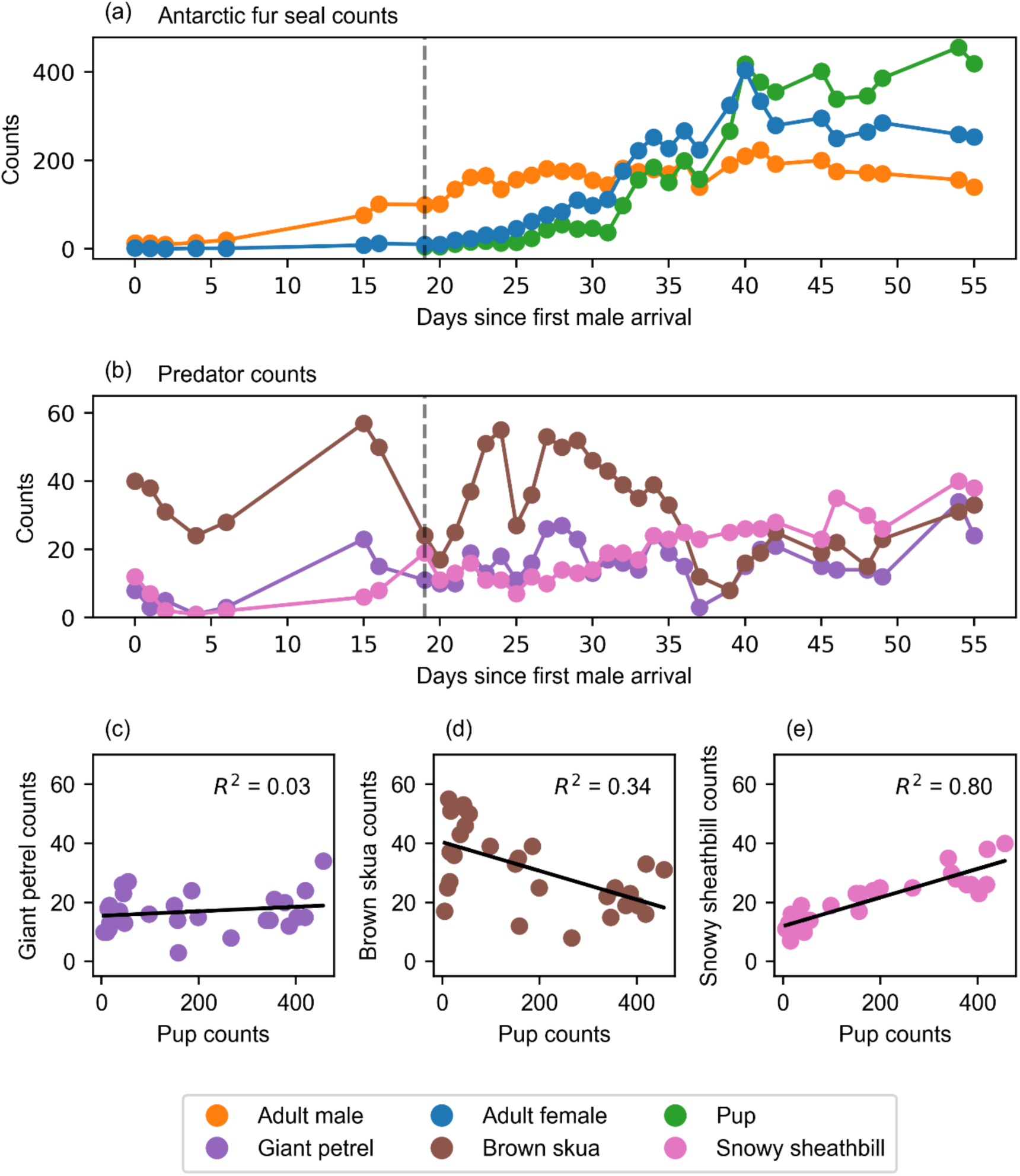
(a) Daily maximum neural network counts of Antarctic fur seals classified as adult males (orange), adult females (blue), and pups (green). (b) Daily maximum counts of avian predator species: giant petrels (purple), brown skuas (brown), and snowy sheathbills (pink). The x-axes in panels (a) and (b) represent days since 30^th^ October 2020, when the first adult male arrived in the colony (defined as ‘day zero’). The vertical black dashed lines in panels (a) and (b) indicate the date of the first-born pup. Panels (c), (d) and (e) show the linear regressions (black lines with corresponding *R²* values) between the maximum daily number of fur seal pups and the maximum daily numbers of (c) giant petrels, (d) brown skuas, and (e) snowy sheathbills.

**Figure 3:**
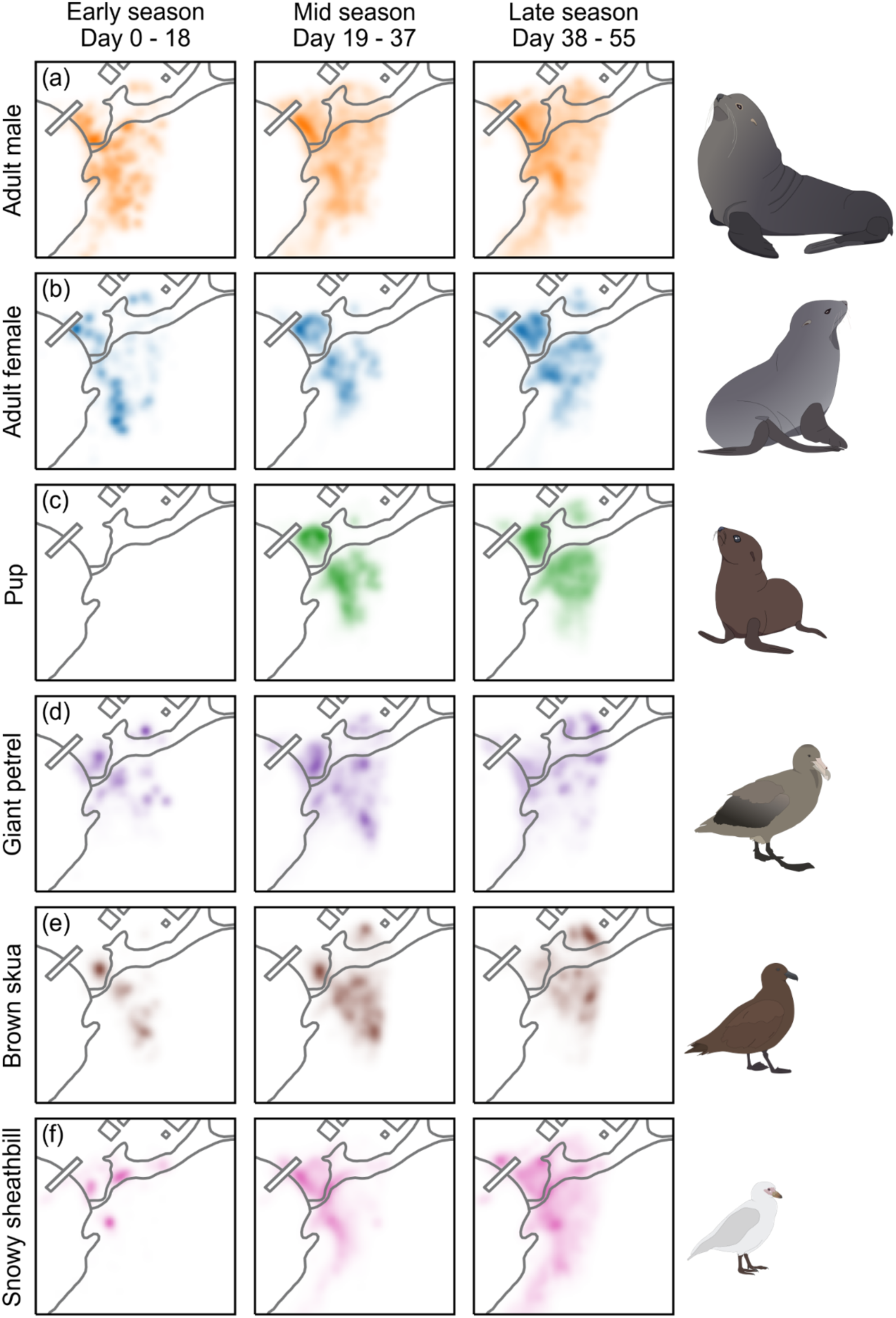
Kernel density estimates for animal detections at freshwater beach, grouped by class and separated into three time periods: Early season (30. Oct to 17. Nov, 19 days), mid-season (18. Nov to 6. Dec, 19 days) and late season (7. Dec to 24. Dec, 18 days). Rows represent detections for each class with original artwork by ALB on the right.

### (e) Data filtering

Daily fluctuations in light conditions significantly influenced visibility. Therefore, we truncated the dataset to include only images captured between 09:00 and 17:00 local time. Additional factors such as fog or water droplets accumulating on the camera lens also reduced image clarity, limited the observable distance, and compromised the field of view. To distinguish usable images for automatic evaluation, the subset of images used for training was manually classified into two categories based on visibility and field of view obstruction. To quantify image quality, we calculated the Laplacian variance (40, see SI Figure 4) for each image using OpenCV (41). The classification threshold was optimized to maximize the F1 score, allowing us to systematically filter out images with insufficient visibility across the entire dataset. Out of a total of 66,645 images, we selected 10,046 (15%) high-quality images that met the criteria of good visibility, sufficient daylight and a clear and unobstructed lens.

**Figure 4:**
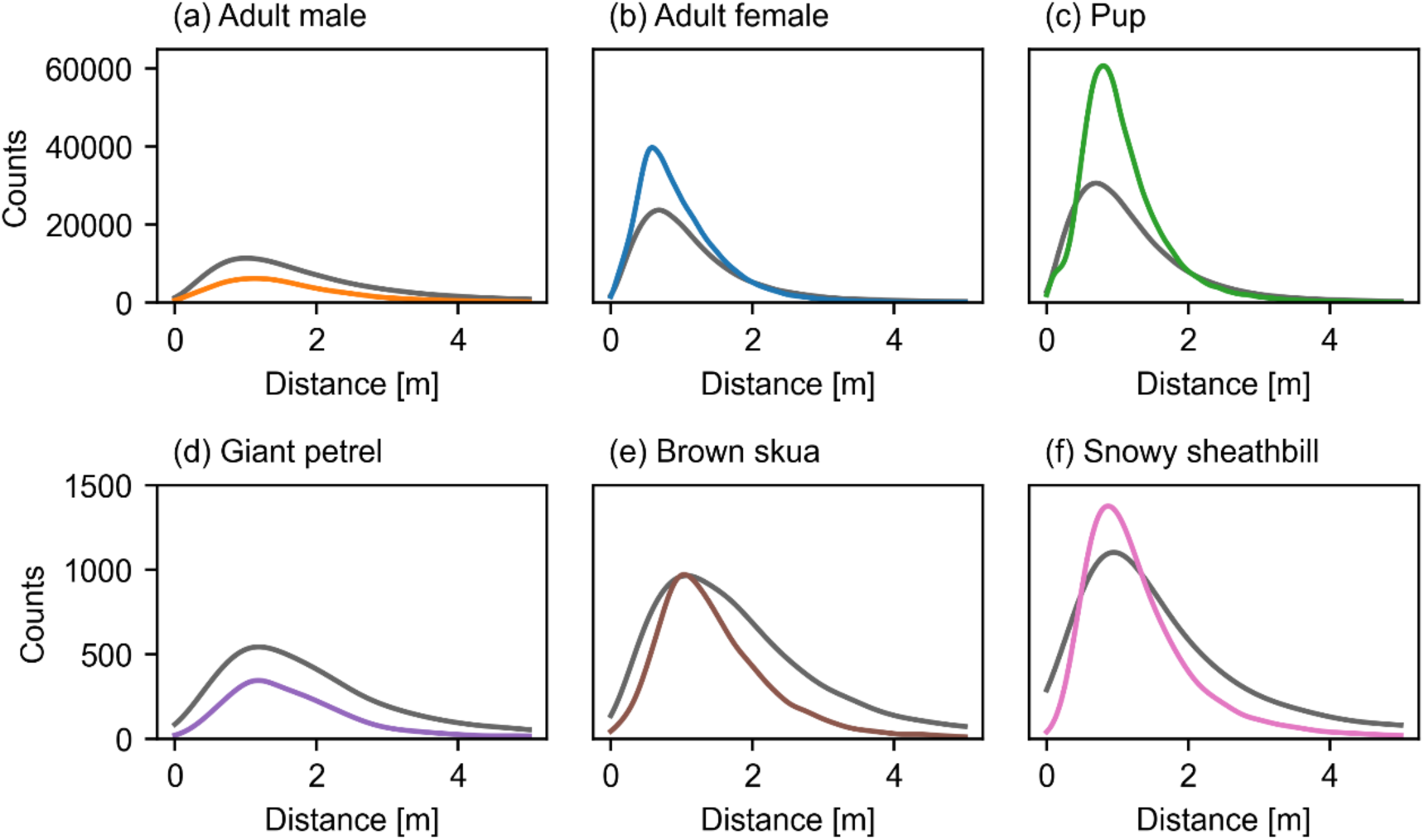
Spatial associations between Antarctic fur seal pups and the six focal classes. Each panel compares the observed and randomized distributions of pup-neighbour distances for different classes of neighbour (adult male, adult female, pup, giant petrel, brown skua and snowy sheathbill). The empirical distributions (coloured lines) were derived from actual pup positions, whereas the randomized distributions (dark grey lines) were generated by introducing a new pup position sampled from a random image, which preserved the global colony structure while disrupting local spatial relationships.

Images only provide a two-dimensional projected view of a three-dimensional scene, so that the distance between the camera and a given individual and the distances between pairs of individuals are not inherently known. We therefore calculated the three-dimensional positions of all observations in space from the image coordinate system using geo-references of the camera position and several landmarks within the camera’s field of view. We referenced our images to the global latitude and longitude coordinate system by manually annotating the positions of landmarks with known positions, and optimizing the camera viewpoint until the projection of space landmark coordinates into the image matched our annotations. For this, we used the Python module CameraTransform (42). Every time the camera alignment changed during the recording period due to maintenance or heavy winds, this procedure was repeated.

To define the geographical boundaries of the Antarctic fur seal colony, we applied kernel density estimation. Specifically, we identified the region containing 99% of all manually annotated Antarctic fur seal detections. By using these manual annotations as a filtering criterion and treating them as ground truth, we excluded automated detections beyond distances where a human observer could no longer reliably classify the animals. This approach ensured that only areas where Antarctic fur seals are most frequently present were analysed, effectively excluding predators that are too far away from Antarctic fur seals pups to pose an imminent threat. Additionally, the BAS field base features a path leading to a jetty, with wooden crates placed alongside it. We did not consider the path, jetty or the crates to be part of the colony. Consequently, all detections within this area were excluded. The excluded detections primarily consisted of snowy sheathbills, which use the crates as a resting spot. Finally, we manually identified the first-born Antarctic fur seal pup in our images (18th Nov) and filtered out pup annotations prior to this date (0.04% of all automatic detections). In addition, we excluded observations of leopard seals, elephant seals, kelp gulls and king penguins due to their rarity. The resulting automated dataset consists of 4.1 million annotated individuals across the breeding season.

### (f) Data analyses

We validated the reliability of the automated counts generated by a neural network by comparing them to the manual counts, revealing a high degree of concordance (see SI Figures 1 and 2). To estimate the abundance of Antarctic fur seal categories (adult males, adult females and pups) and the avian species (giant petrel, brown skua and snowy sheathbill) throughout the season, we used the daily maximum count for each observed class. We tested for associations between the numbers of birds and pups by fitting a linear regression model separately for each bird species, and we used the corresponding *R*² values to quantify the strength of the associations.

To characterize the spatial distribution of the animals and to identify their habitat preferences, we applied kernel density estimation to each class. To explore temporal changes, we divided the recording period into three equal time intervals (early season: 30^th^ Oct till 17^th^ Nov, mid-season: 18^th^ Oct till 6^th^ Dec, late season: 7^th^ Dec till 24^th^ Dec), with the birth of the first pup marking the start of the mid-season.

Fine-scale spatial associations between Antarctic fur seal pups and the focal classes were investigated by comparing the distributions of observed and random geographical distances among individuals. For each pup, we calculated the distance to its nearest neighbour and grouped them based on the neighbour’s class. To generate the null expectation, we implemented a randomized pup placement method that preserved the overall spatial arrangement of pups across the season but disrupted local spatial relationships. Specifically, for each pup in every image, we randomly selected the position of a pup from another image and calculated nearest neighbour distances in the same way as for the observed pup. Deviations between observed and random distance distributions indicate local attraction or repulsion between pups and other classes.

To investigate fine-scale predator-prey interactions, we analysed spatial relationships among the three Antarctic fur seal categories and the avian species. For each bird, we calculated the Euclidean distance to all pups within a radius of 10m. Pup proximity to a predator was used as a proxy for predation risk, assuming that pups can mitigate this risk by staying close to an adult Antarctic fur seal. Accordingly, for each pup, we also recorded the distance to its nearest Antarctic fur seal neighbour.

### (g) Data availability

All data supporting the findings of this study—including the trained model, annotated training images, and the resulting manual and automated counts—are available in the Zenodo repository (DOI: 10.5281/zenodo.18141470). The code required to reproduce this study, including Jupyter notebooks for neural network training and inference, as well as for the entire analysis are available on GitHub (https://github.com/fabrylab/AntarcticFurSealPredatorPrey). All analyses were conducted using Python.

## 3. Results

Our neutral network identified a total of 4.1 million detections across six focal classes (Antarctic fur seal categories: adult male, adult female, pup; avian species: giant petrel, brown skua and snowy sheathbill, Figure 1c, SI Figures 5 and 6) from images collected at FWB (Figure 1b) at one-minute intervals.

**Figure 5:**
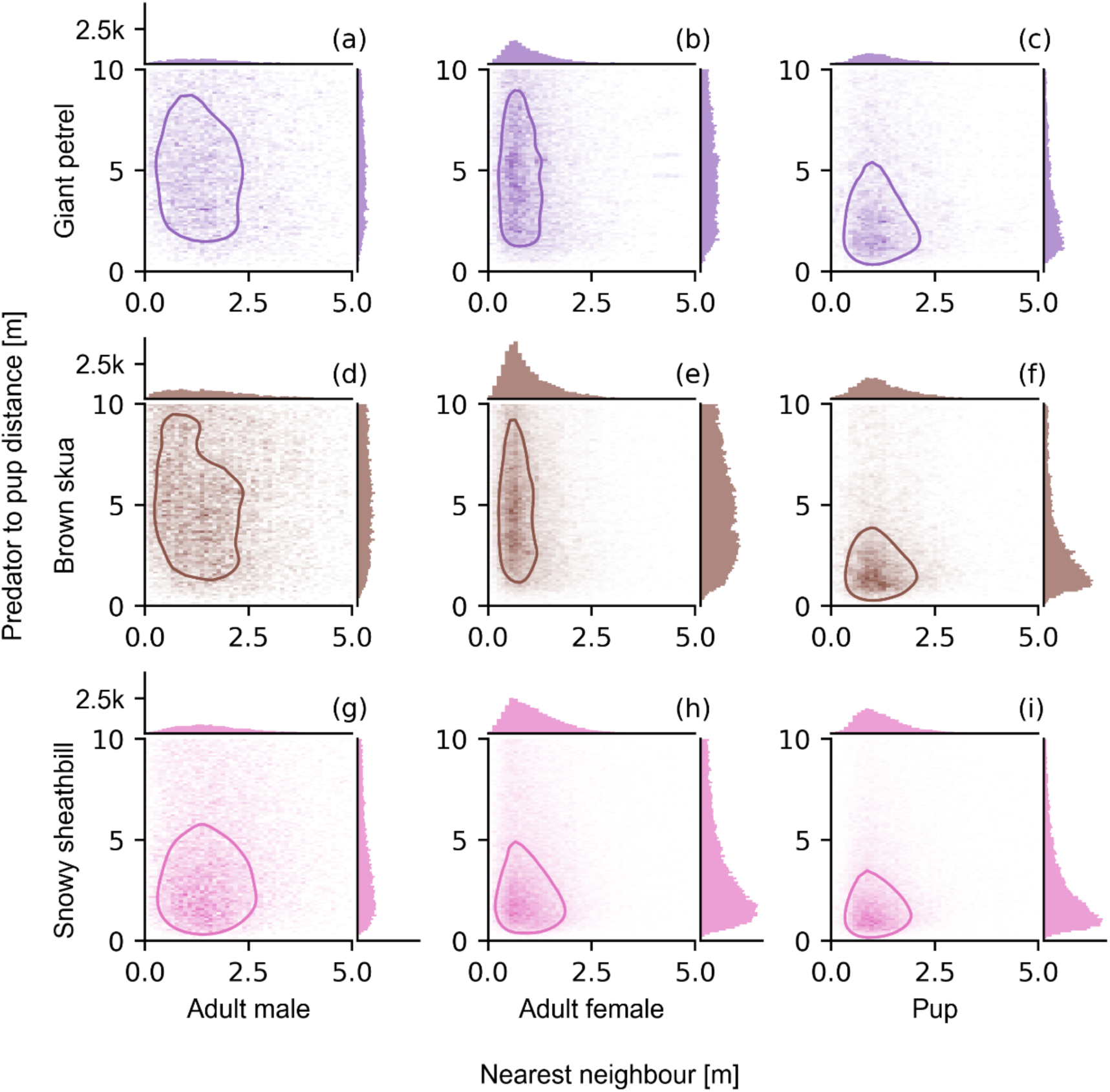
Spatial relationships between Antarctic fur seal pups and their avian predators, and the influence of neighbouring Antarctic fur seals on predation risk. Each panel shows a 2D-histogram depicting the distance between a predator and a pup within a 10-meter radius (y axis) versus the distance from that same pup to its nearest neighbour (x axis). Also shown are the 50% density contour lines. The top row (panels a,b,c) shows the histograms for giant petrels, the middle row (panels d, e, f) for brown skuas, and the bottom row (panels g, h, i) for snowy sheathbills. The left column (panels a, d, g) shows the histograms where the nearest neighbour of the prey pup was an adult male, the middle column (panels b, e, h) where the nearest neighbour of the prey pup was an adult female, and the right column (panels c, f, i) where the nearest neighbour of the prey pup was another pup. Shown alongside each panel are 1D-histograms depicting distributions of predator-to-prey distances (right), and prey-to-nearest neighbour distances (top). The histogram bin width is 0.1 m and histogram height (indicating the number of observations per bin) is scaled as indicated.

### (a) Temporal patterns of abundance

To analyse temporal patterns of abundance, we plotted the daily maximum counts of each focal class over the course of the breeding season, starting on October 30th and ending on December 24th for a total of 56 days (Figure 2 a, b). Focusing initially on Antarctic fur seals, only adult males were present in the first two weeks and their numbers gradually increased until day 30, after which they remained relatively stable. Adult females began arriving at the colony around day 15, their numbers peaked on day 40 and thereafter declined. The first pup was born on day 19, after which the number of pups steadily increased. These temporal patterns align with the known breeding phenology of Antarctic fur seals (24).

The number of giant petrels fluctuated throughout the season (daily count across all images = 16 ± 7 birds) without any clear temporal pattern. Brown skuas were even more variable, peaking during the first four weeks of the season and declining after day 27. By contrast, snowy sheathbills showed smaller daily fluctuations and gradually increased in abundance over the season, from 12 individuals in a single image on day zero to 38 on the last day, with a maximum of 41 on the 23^rd^ of December.

To investigate temporal associations between pups and the avian predator-scavenger species, we regressed daily maximum bird counts against the daily maximum pup counts. There was no association between the abundance of pups and giant petrels (*R²* = 0.03, *p* = 0.40; Figure 2c). Brown skuas exhibited a significant but weak negative association (*R²* = 0.33, *p* = 0.0014; Figure 2d), whereas snowy sheathbills exhibited a strong positive association (*R²* = 0.80, *p* < 0.001; Figure 2e), suggesting a direct link between the abundance of pups and sheathbills.

### (b) Spatial patterns of abundance

Next, we analysed the spatial distribution of the Antarctic fur seal categories and avian species across three consecutive time periods (Figure 3): the early season (day 0–18) when no pups were present; the mid-season (day 19–37) when pup and adult female counts steadily increased, and the late season (after day 38) when the number of adult females began to decline. The patchy distributions observed during the first 19 days for all classes can be partially attributed to the small number of days with good visibility (seven in total). Across the season, adult males occupied the largest area, with a tendency to cluster near the shoreline, and their distribution remained relatively stable over time. Adult females were more concentrated in central areas of the colony, except during the early season, when their distribution overlapped more with the adult males. Pups were predominantly found in areas with a high density of adult females, reflecting their close association. Adult females and pups, but not adult males, avoided the freshwater stream, suggesting that it represents lower-quality habitat mainly occupied by less competitive adult males.

Although spatial distributions of the three avian species overlapped to some extent with the Antarctic fur seal colony, their spatial associations differed markedly. Only giant petrels were found in the tidal zone and shallow water. Their distribution was similar throughout the season and was not clearly associated with any of the Antarctic fur seal categories. By contrast, the spatial distribution of brown skuas closely matched that of adult female Antarctic fur seals and their pups, especially during the mid and late season, suggesting a potential ecological interaction. Snowy sheathbills, on the other hand, showed an expanding geographical distribution over time, reflecting their steady increase in numbers.

### (c) Fine-scale associations between pups and their nearest neighbours

We found that pups were more likely to be positioned near other pups or adult females in the observed dataset compared to the random dataset, indicating spatial attraction (Figure 4). Conversely, fewer pups were observed in close proximity to adult males, giant petrels or brown skuas, implying spatial avoidance. Similarly, while pups were less frequently observed near snowy sheathbills, for those that were, their observed distances were smaller than expected by chance.

### (d) Fine-scale predator-prey interactions

The results of the nearest-neighbour analysis are shown in nine two-dimensional (bivariate) histograms (Figure 5). These histograms illustrate relationships between pup-to-predator distances and pup-to-nearest Antarctic fur seal distances. The histograms are arranged in a 3 x 3 grid, with the rows representing the gradient of avian predators to scavengers (giant petrels, brown skuas and snowy sheathbills) and the columns representing the three classes of Antarctic fur seal neighbour (adult male, adult female and pup). Darker shades indicate more observations, and 50% isopleth lines enclose the densest 50% of all observations. To complement these visualisations, each 2D histogram is accompanied by one-dimensional marginal histograms of pup-neighbour distances on the x-axis and pup-bird distances on the y-axis. Narrow distributions in these plots could indicate either attraction or avoidance between the avian predator and a specific pup-Antarctic fur seal neighbour pair.

Giant petrels and brown skuas maintained the largest distances from pups accompanied by adult males (left column of figure 5, the nearest neighbour is an adult male Antarctic fur seal, median distance = 1.3 m apart), with distances of around 3.2 m for giant petrels, 4.5 m for brown skuas, and 1.8 m for snowy sheathbills. Similarly large distances to predators were observed when the nearest neighbour of the pups was an adult female Antarctic fur seal (middle column of figure 5, the nearest neighbour is an adult female, median distance = 0.6 m), with distances of around 3.7 m for giant petrels and 3 m for brown skuas, whereas snowy sheathbills kept a distance of only 1.5 m. This indicates that pups in close proximity to adult Antarctic fur seals of both sexes were avoided by brown skuas and giant petrels, but not by the snowy sheathbills. This pattern may reflect the tendency of adult Antarctic fur seals to deter or chase away potential predators, particularly giant petrels and brown skuas which pose a greater predation threat.

All three bird species closely approached unaccompanied pups (left column, the nearest neighbour is another pup, median distance = 1 m apart), with median distances of around 1.5 m for giant petrels, 1.3 m for brown skuas, and 1.1 m for snowy sheathbills. Furthermore, the 2D distributions were right skewed, as indicated by asymmetry in the isopleth lines, with more observations at closer avian predator-pup distances and larger pup-pup distances. This suggests that isolated pups are more vulnerable to predation than those occupying higher density areas. The 1-D histograms of pup-neighbour distances (above each 2-D histogram, figure 5) further indicate that adult female Antarctic fur seals were the most frequent nearest neighbours of pups, reflecting nutritional dependence and the protective role of mothers. By contrast, pups tended to avoid adult male Antarctic fur seals despite the potential protection they may provide from predators (Figure 5, left column). As traumatic injuries caused by territorial males are a major cause of early-life mortality (13), this pattern suggests a trade-off: pups may benefit from staying close to adults for protection but must maintain a safe distance from adult males to avoid accidental trampling.

## 4. Discussion

### Automated camera system and neural network analysis

The automated daily maximum counts for all three avian predators consistently exceeded the manual counts (SI Figure 5 and 6). This discrepancy arises because the manual annotations were based on a single snapshot per day, whereas the automated detection system evaluated all of the available images (up to 512 per day under optimal weather conditions), making it unlikely that a single image captured the peak number of avian predators on a given day. Daily fluctuations in avian predator numbers are also expected as these species move back and forth between nesting and foraging sites throughout the day (16,19,43). By contrast, male Antarctic fur seal hold territories for an average of 30 days (44) and exhibit strong site fidelity (45), resulting in relatively stable counts, while adult females give birth and often mate before their first postpartum foraging trip (24), resulting in predictable periods of presence and absence. Ultimately, the strong relationship between the manual and neural network counts (F1 = 0.75) demonstrates that the automated detections are reliable.

### Temporal patterns of abundance

To investigate temporal abundance patterns, we plotted daily maximum neural network counts separately for each focal class. The observed patterns in Antarctic fur seal abundance closely matched the species’ known breeding phenology, with males arriving about a month before the females (23), which give birth shortly after coming ashore, suckle their pups for around a week, and then alternate foraging trips at sea with provisioning bouts ashore (24). This concordance between our results and established patterns further validates the automated detection approach.

If Antarctic fur seal pups serve as an important food resource for avian predators, we would expect to find strong temporal associations between pup and predator abundance. Giant petrel counts remained relatively stable throughout the season and accordingly, no association was found with pup abundance. This may reflect intraspecific competition, as giant petrels are known to exhibit strong antagonistic behaviour that may limit the number of giant petrels that can forage within a given area (46,47). Surprisingly, we observed a weak negative temporal association between the abundance of pups and brown skuas, despite previous reports indicating that brown skuas rely on fur seal resources throughout their breeding season (16). Brown skua numbers declined around two thirds of the way through the study period, coinciding with the hatching period of their chicks (16). While seal carrion dominates the diet of brown skuas during incubation, they shift to seabird prey during chick-rearing, leaving only non-breeding adults feeding on seal resources, which may explain our findings (16,48). By contrast, the strong positive temporal association between snowy sheathbill abundance and pup numbers likely reflects the birds’ opportunistic foraging on placentae, carrion and faeces, which are particularly abundant at the height of the fur seal breeding season (19).

### Spatial patterns of abundance

Differences in the spatial distributions of the three avian species likely indicate foraging niche partitioning. Giant petrels prey on Antarctic fur seal pups using two distinct techniques: pecking and drowning. The latter involves dragging pups into shallow water and blocking their return to land (15), which may explain why giant petrels, unlike the other avian predators, extend their spatial distribution into the shallow water. Brown skuas exhibited strong spatial overlap with adult female Antarctic fur seals and their pups, likely reflecting their dependence on placentae (16). By contrast, snowy sheathbills occupied a broader area, likely reflecting their more generalist scavenging behaviour (19) and / or physical displacement by the other predators.

### Fine-scale pup association patterns

We investigated fine-scale associations between Antarctic fur seal pups and the focal classes by comparing the empirical distributions of pup-to-nearest-neighbour distances with randomized distributions. A greater than expected number of pups in close proximity to a given class indicates local attraction, whereas fewer than expected observations indicates avoidance. As anticipated, given that our study coincided with a period when Antarctic fur seal pups are nutritionally dependent on their mothers, we found strong evidence of local attraction between pups and adult females (49). This observation aligns with previous research suggesting that pups near their mothers can more easily fend off predators than unattended pups (15). Additionally, we observed local attraction among pups, which could facilitate social interactions while their mothers are away from the colony foraging at sea (50). Alternatively, the grouping together of pups could serve as an anti-predator response, as this can increase vigilance and improve predator detection (51).

Conversely, Antarctic fur seal pups appear to actively avoid both avian predators and adult male Antarctic fur seals. This avoidance was most pronounced for adult male Antarctic fur seals, historically the leading cause of pup mortality (18), and for giant petrels, which are generally considered their main predator (15), highlighting the risks posed by these two classes. Pups also avoided brown skuas and, to a lesser extent, snowy sheathbills. This gradient of avoidance appears to be linked to predator size, potentially reflecting the relative level of threat posed by each species, decreasing from giant petrels through brown skuas to snowy sheathbills. This pattern aligns with the threat-sensitivity hypothesis, which predicts that prey animals should adjust their antipredator behaviour according to perceived risk (52). Evidence supporting this hypothesis has been found in other taxa (53–55) and juveniles of both Cape fur seals (*Arctocephalus pusillus pusillus*) and Antarctic fur seals have been observed to adjust their movement patterns under high predation risk (50,56). Taken together, these results suggest that pups may be capable of assessing and responding to varying levels of predation risk, a hypothesis that could be tested experimentally by presenting them with models of predators differing in size and morphology.

### Fine-scale predator-prey interactions

We implemented a nearest-neighbour analysis to investigate fine-scale predator-prey interactions, complementing the fine-scale pup association patterns by taking the perspective of individual predators. Although our remote observation approach allowed accurate, continuous monitoring of animal positions, its low frame rate (one image per minute) limited our ability to differentiate between scavenging and active predation events. Consequently, we used proximity to predators as a proxy for pup predation risk. Pups whose nearest neighbour was another pup had predators closer to them, consistent with previous observations that unattended pups are particularly vulnerable to predation (15).

While fine-scale pup association patterns showed that Antarctic fur seal pups avoid adult males, the presence of both adult males and females was associated with reduced predation risk from giant petrels and brown skuas, highlighting the critical role of adult females in mitigating predation risk. By contrast, the consistent spatial avoidance of adult males by pups, giant petrels and brown skuas suggests that any protective effect of adult males is likely incidental rather than intentional. Notably, snowy sheathbills were observed near pups regardless of the presence of adult seals, reflecting higher tolerance of this species by adult fur seals or *vice versa*. Although rare instances of sheathbill predation on pups have been documented (18) these remarkably resourceful birds exploit a broad range of available resources, including blood and even milk from southern elephant seals (20), and are principally opportunistic scavengers (19,20). Their preference for scavenging, along with the incidental hygienic benefits it may provide, likely explains the higher tolerance of their presence near Antarctic fur seal pups.

Overall, our study provides detailed insights into spatiotemporal associations and predator-prey interactions in a breeding colony of Antarctic fur seals. It highlights the trade-offs that fur seal pups must navigate between nutritional dependence on their mothers, social development, and the need to avoid avian predators and territorial male fur seals. Additionally, our findings reveal clear spatial niche partitioning within a guild of predator-scavengers, where giant petrels control the shoreline and, together with brown skuas, predominantly occupy central areas of the colony, whereas snowy sheathbills range throughout the entire area. Further research could explore the possibility of a hierarchical organisation within this guild in which giant petrels, by virtue of their size and aggressive behaviour, monopolize the highest quality resources (46), followed by brown skuas, with snowy sheathbills relegated to scavenging the leftovers.

### (f) Caveats

Investigating predator-prey interactions in natural systems is inherently complex and often requires the use of proxies such as spatial overlap to infer predation risk (57). In our study, the fixed camera captured a single image per minute, limiting our ability to directly observe predation events, which would be the ideal metric for investigating predator-prey interactions. Instead, we used temporal and spatial overlap as initial proxies, followed by a fine-scale analysis of the distances between Antarctic fur seal pups and avian predators. While computer-based visual detections are less precise than manual annotations, the large volume of data they generate greatly increases statistical power, which would have been difficult to achieve otherwise.

Nevertheless, using a fixed camera set-up has certain limitations. First, adverse weather conditions produced some gaps in our dataset, particularly early in the season. However, we believe these had a minimal impact on our analysis, as the presence of animals was generally low during this period and our predator–prey interaction data were restricted to the period following the birth of the first pup. Second, our observations were limited to daylight hours, since we deployed a standard camera. In future studies, this issue could be resolved by deploying an additional infrared camera. Third, our observations were constrained by the camera’s field of view. Therefore, objects appeared smaller and at lower resolution with increasing distance from the camera, limiting our ability to detect objects in peripheral areas of the colony. Furthermore, pups transition into the tussock grass behind the colony as they develop better motor skills, which is a potential predator avoidance behaviour (50). This seasonal shift limits our ability to observe the pups as they approach weaning, although most pup mortalities occur before this transition (49,58).

### Future perspectives

Hardware improvements to the current camera setup could significantly expand research opportunities and enable the investigation of a wider range of ecological questions. Establishing a permanent power supply and data connection at the study site would greatly reduce maintenance demands and facilitate multi-year monitoring. Long-term observations are essential for detecting broader ecological trends and monitoring changes in population dynamics and habitat use. Increasing the camera frame rate would further allow more detailed investigation of predation events as well as interactions among predators. A permanent camera installation would therefore complement traditional field-based approaches by providing continuous, unobtrusive data collection. Importantly, the neural network trained in this study can be readily reused for future monitoring efforts, requiring only minimal retraining to adapt to subsequent years or different locations. This transferability reduces the amount of data needed for adaptation, making this approach both scalable and cost-effective for long-term monitoring of predator-prey interactions and population trends in remote ecosystems.

More broadly, automated camera systems offer a non-invasive, labour-efficient means of gaining insights into animal behaviour and ecology. Their popularity has grown substantially over the past decade, driven by improved availability and advances in machine-learning algorithms capable of processing large datasets (59). The approach presented here could be readily applied to other colonial species, and similar methods have been used to estimate density in colonies containing multiple species (60) and to monitor species interactions underwater (61). Collectively, these advances highlight the growing role of automated imagery as a complementary approach to traditional methods, especially in remote or challenging environments.

## Conclusion

Despite the limitations of our study, the use of a low-cost, automated camera system coupled with neural network-based analysis proved highly effective at generating a large, high-quality spatiotemporal dataset simultaneously covering multiple species. This approach offers significant advantages over traditional field methods by minimising human disturbance and enabling robust statistical inference of spatial and temporal patterns. Compared to classical field observations, it offers comprehensive and near-continuous coverage, facilitating the detection of fine-scale dynamics that might otherwise go unnoticed. Applying this approach to an Antarctic fur seal breeding colony allowed us to characterise fine-scale spatiotemporal patterns and gain fresh insights into fine-scale predator-prey interactions.

## Acknowledgements

The authors thank Iain Angus Gordon and David Reid for technical support in the field. The authors gratefully acknowledge the scientific support and HPC resources provided by the Erlangen National High Performance Computing Center (NHR@FAU) of the Friedrich-Alexander-Universität Erlangen-Nürnberg (FAU). Support for the article processing charge was granted by the DFG and the Open Access Publication Fund of Bielefeld University.

## Funding Statement

A.L.B. Collaborative Research Centre Transregio 212 “A Novel Synthesis of Individualisation across Behaviour, Ecology and Evolution: Niche Choice, Niche Conformance, Niche Construction (NC3)” (SFB TRR 212, project numbers 316099922 & 396774617)

J.B. DFG (German Research Foundation) priority programme “Antarctic Research with Comparative Investigations in Arctic Ice Areas” (SPP 1158, project number ZI1527/7-1)

A.W. DFG (German Research Foundation) priority programme “Antarctic Research with Comparative Investigations in Arctic Ice Areas” (SPP 1158, project number 443134677) C.F.-C. The core science programme of the British Antarctic Survey, Polar Science For a Sustainable Planet, NERC-UKRI (Natural Environment Research Council, United Kingdom UKRI).

J.F. The core science programme of the British Antarctic Survey, Polar Science For a Sustainable Planet, NERC-UKRI (Natural Environment Research Council, United Kingdom UKRI).

R.N. Collaborative Research Centre Transregio 212 “A Novel Synthesis of Individualisation across Behaviour, Ecology and Evolution: Niche Choice, Niche Conformance, Niche Construction (NC3)” (SFB TRR 212, project numbers 316099922 & 396774617)

B.F. DFG (German Research Foundation) priority programme “Antarctic Research with Comparative Investigations in Arctic Ice Areas” (SPP 1158, project number 443134677)

J.I.H. DFG (German Research Foundation) priority programme “Antarctic Research with Comparative Investigations in Arctic Ice Areas” SPP 1158 (project number 424119118). Collaborative Research Centre Transregio 212 “A Novel Synthesis of Individualisation across Behaviour, Ecology and Evolution: Niche Choice, Niche Conformance, Niche Construction (NC3)” (SFB TRR 212, project numbers 316099922 & 396774617).

## Ethical Statement

The work at Bird Island was carried out by BAS under permits from the Government of South Georgia and the South Sandwich Islands (Wildlife and Protected Areas Ordinance (2011), RAP permit number 2020/015).

## Data Accessibility

The manual and automated counts, positions, classes are available on GitHub (https://github.com/fabrylab/AntarcticFurSealPredatorPrey). All software necessary to reproduce the study is publicly available in the repository. All data supporting the finding of this study – including the trained model, annotated training images and the resulting manual and automated counts – are available on Zenodo (DOI: 10.5281/zenodo.18141470).

## Authors’ Contributions

J.I.H., R.N. and B.F. conceived the study. J.I.H. acquired funding for the project. R.N. and C.F.-C. conducted the fieldwork and C.F.-C. maintained the camera. A.W. trained the neural network. J.B. in discussion with A.L.B. and B.F. performed the formal analyses and visualized the results.

A.L.B., A.W., J.B., B.F. and J.I.H. wrote the original draft. All authors provided feedback and approved the final manuscript.

## References

1. Hastings A. Complex Interactions Between Dispersal and Dynamics: Lessons From Coupled Logistic Equations. Ecology. 1993;74(5):1362–72.

2. Reid K, Croxall JP. Environmental response of upper trophic-level predators reveals a system change in an Antarctic marine ecosystem. Proc Biol Sci. 2001 Feb 22;268(1465):377–84.

3. Spraker TR, DeLong RL, Lyons ET, Melin SR. HOOKWORM ENTERITIS WITH BACTEREMIA IN CALIFORNIA SEA LION PUPS ON SAN MIGUEL ISLAND. J Wildl Dis. 2007 Apr 1;43(2):179–88.

4. Boveng PL, Hiruki LM, Schwartz MK, Bengtson JL. Population Growth of Antarctic Fur Seals: Limitation by a Top Predator, the Leopard Seal? Ecology. 1998;79(8):2863–77.

5. Allee WC (Warder Clyde). Animal aggregations, a study in general sociology [Internet]. Chicago, The University of Chicago Press, 1931; 1931. 452 p. Available from: https://www.biodiversitylibrary.org/item/31040

6. Kramer AM, Dennis B, Liebhold AM, Drake JM. The evidence for Allee effects. Popul Ecol. 2009;51(3):341.

7. Bednekoff PA, Lima SL. Re–examining safety in numbers: interactions between risk dilution and collective detection depend upon predator targeting behaviour. Proc R Soc Lond B Biol Sci. 1998 Oct 22;65(1409):2021–6.

8. Foster WA, Treherne JE. Evidence for the dilution effect in the selfish herd from fish predation on a marine insect. Nature. 1981 Oct 1;293:466–7.

9. Stephens PA, Sutherland WJ. Consequences of the Allee effect for behaviour, ecology and conservation. Trends Ecol Evol. 1999 Oct 1;14(10):401–5.

10. Nagel R, Stainfield C, Fox-Clarke C, Toscani C, Forcada J, Hoffman J. Evidence for an Allee effect in a declining fur seal population [Internet]. Dryad; 2021 [cited 2022 Oct 6]. p. 92074 bytes. Available from: http://datadryad.org/stash/dataset/doi:10.5061/dryad.zcrjdfnb0

11. Doidge DW, Croxall JP, Baker JR. Density-dependent pup mortality in the Antarctic fur seal Arctocephalus gazellu at South Georgia. J Zool. 1984;202(3):449–60.

12. Hunt G, Heinemann D, Everson I. Distributions and predator-prey interactions of macaroni penguins, Antarctic fur seals, and Antarctic krill near Bird Island, South Georgia. Mar Ecol Prog Ser. 1992;86:15–30.

13. Reid K, Forcada J. Causes of offspring mortality in the Antarctic fur seal, *Arctocephalus gazella*: the interaction of density dependence and ecosystem variability. Can J Zool. 2005 Apr 1;83(4):604–9.

14. Forcada J, Hoffman JI. Climate change selects for heterozygosity in a declining fur seal population. Nature. 2014 Jul;511(7510):462–5.

15. Nagel R, Coleman J, Stainfield C, Forcada J, Hoffman JI. Observations of Giant Petrels (Macronectes sp.) Attacking and Killing Antarctic Fur Seal (Arctocephalus gazella) Pups. Aquat Mamm. 2022 Nov 15;48(6):509–12.

16. Phillips RA, Phalan B, Forster IP. Diet and long-term changes in population size and productivity of brown skuas Catharacta antarctica lonnbergi at Bird Island, South Georgia. Polar Biol. 2004 Aug 1;27(9):555–61.

17. González-Solís J, Croxall JP, Wood AG. Foraging partitioning between giant petrels Macronectes spp. and its relationship with breeding population changes at Bird Island, South Georgia. Mar Ecol Prog Ser. 2000 Oct 5;204:279–88.

18. Baker JR, Doidge DW. Pathology of the antarctic fur seal (Arctocephalus gazella) in South Georgia. Br Vet J. 1984 Mar;140(2):210–9.

19. Burger AE. Time Budgets, Energy Needs and Kleptoparasitism in Breeding Lesser Sheathbills (Chionis minor). The Condor. 1981 May;83(2):106–12.

20. Favero M. Foraging Ecology of Pale-Faced Sheathbills in Colonies of Southern Elephant Seals at King George Island, Antarctica (La Ecología de la Alimentación de la Chionis alba en los Harenes de Elefantes Marinos en la Isla King George, Antarctica). J Field Ornithol. 1996;67(2):292–9.

21. Mattisson J, Rauset GR, Odden J, Andrén H, Linnell JDC, Persson J. Predation or scavenging? Prey body condition influences decision-making in a facultative predator, the wolverine. Ecosphere. 2016;7(8):e01407.

22. Wilson EE, Wolkovich EM. Scavenging: how carnivores and carrion structure communities. Trends Ecol Evol. 2011 Mar 1;26(3):129–35.

23. McCann TS. Territoriality and breeding behaviour of adult male Antarctic Fur seal, Arctocephalus gazella. J Zool. 1980;192(3):295–310.

24. Duck CD. Annual variation in the timing of reproduction in Antarctic fur seals, Arctocephalus gazella, at Bird Island, South Georgia. J Zool. 1990;222(1):103–16.

25. Lunn NJ, Boyd IL, Croxall JP. Reproductive Performance of Female Antarctic Fur Seals: The Influence of Age, Breeding Experience, Environmental Variation and Individual Quality. J Anim Ecol. 1994;63(4):827–40.

26. Hamilton WD. Geometry for the selfish herd. J Theor Biol. 1971 May 1;31(2):295–311.

27. Krause J. Differential fitness returns in relation to spatial position in groups. Biol Rev Camb Philos Soc. 1994 May;69(2):187–206.

28. Forcada J, Hoffman JI, Gimenez O, Staniland IJ, Bucktrout P, Wood AG. Ninety years of change, from commercial extinction to recovery, range expansion and decline for Antarctic fur seals at South Georgia. Glob Change Biol [Internet]. 2023 Oct 15 [cited 2023 Oct 19];n/a(n/a). Available from: https://onlinelibrary.wiley.com/doi/abs/10.1111/gcb.16947

29. McMahon CR, Howe H, Hoff J van den, Alderman R, Brolsma H, Hindell MA. Satellites, the All-Seeing Eyes in the Sky: Counting Elephant Seals from Space. PLOS ONE. 2014 Mar 20;9(3):e92613.

30. LaRue MA, Rotella JJ, Garrott RA, Siniff DB, Ainley DG, Stauffer GE, et al. Satellite imagery can be used to detect variation in abundance of Weddell seals (Leptonychotes weddellii) in Erebus Bay, Antarctica. Polar Biol. 2011 Nov 1;34(11):1727–37.

31. Carroll D, Infantes E, Pagan EV, Harding KC. Approaching a population-level assessment of body size in pinnipeds using drones, an early warning of environmental degradation. Remote Sens Ecol Conserv [Internet]. 2024 [cited 2025 Jan 10];n/a(n/a). Available from: https://onlinelibrary.wiley.com/doi/abs/10.1002/rse2.413

32. Hoekendijk JPA, Grundlehner A, Brasseur S, Kellenberger B, Tuia D, Aarts G. Stay close, but not too close: aerial image analysis reveals patterns of social distancing in seal colonies. R Soc Open Sci. 2023 Aug 9;10(8):230269.

33. Winterl A, Richter S, Houstin A, Nesterova AP, Bonadonna F, Schneider W, et al. micrObs – A customizable time-lapse camera for ecological studies. HardwareX. 2020 Oct 1;8:e00134.

34. Gerum RC, Richter S, Fabry B, Zitterbart DP. ClickPoints: an expandable toolbox for scientific image annotation and analysis. Methods Ecol Evol. 2017;8(6):750–6.

35. Tan M, Le QV. EfficientNet: Rethinking Model Scaling for Convolutional Neural Networks [Internet]. arXiv; 2020 [cited 2024 Dec 6]. Available from: http://arxiv.org/abs/1905.11946

36. Redmon J, Divvala S, Girshick R, Farhadi A. You Only Look Once: Unified, Real-Time Object Detection [Internet]. arXiv; 2016 [cited 2024 Dec 6]. Available from: http://arxiv.org/abs/1506.02640

37. Abadi M, Barham P, Chen J, Chen Z, Davis A, Dean J, et al. TensorFlow: A system for large-scale machine learning [Internet]. arXiv; 2016 [cited 2024 Dec 6]. Available from: http://arxiv.org/abs/1605.08695

38. Chollet F, others. Keras. 2015.

39. Kingma DP, Ba J. Adam: A Method for Stochastic Optimization [Internet]. arXiv; 2017 [cited 2025 Mar 6]. Available from: http://arxiv.org/abs/1412.6980

40. Pech-Pacheco JL, Cristobal G, Chamorro-Martinez J, Fernandez-Valdivia J. Diatom autofocusing in brightfield microscopy: a comparative study. In: Proceedings 15th International Conference on Pattern Recognition ICPR-2000 [Internet]. 2000 [cited 2025 Feb 19]. p. 314–7 vol.3. Available from: https://ieeexplore.ieee.org/document/903548

41. Bradski G. The OpenCV Library, Dr. Dobb’s Journal of Software Tools. 2000 [cited 2025 Feb 19]; Available from: https://search.hsl.med.nyu.edu

42. Gerum RC, Richter S, Winterl A, Mark C, Fabry B, Le Bohec C, et al. CameraTransform: A Python package for perspective corrections and image mapping. SoftwareX. 2019 Jul 1;10:100333.

43. Poncet S, Wolfaardt AC, Barbraud C, Reyes-Arriagada R, Black A, Powell RB, et al. The distribution, abundance, status and global importance of giant petrels (Macronectes giganteus and M. halli) breeding at South Georgia. Polar Biol. 2020 Jan 1;43(1):17–34.

44. Boyd IL, Duck CD. Mass Changes and Metabolism in Territorial Male Antarctic Fur Seals (Arctocephalus gazella). Physiol Zool. 1991 Jan;64(1):375–92.

45. Hoffman JI, Boyd IL, Amos W. MALE REPRODUCTIVE STRATEGY AND THE IMPORTANCE OF MATERNAL STATUS IN THE ANTARCTIC FUR SEAL ARCTOCEPHALUS GAZELLA. 2003 Feb 20;14.

46. de Bruyn PJN, Cooper J. Who’s the boss? Giant petrel arrival times and interspecific interactions at a seal carcass at sub-Antarctic Marion Island. Polar Biol. 2005 Jun 1;28(7):571–3.

47. Hunter S. The food and feeding ecology of the giant petrels Macronectes halli and M. giganteus at South Georgia. J Zool. 1983;200(4):521–38.

48. Borghello P, Torres DS, Montalti D, Ibañez AE. Diet of the Brown Skua (Stercorarius antarcticus lonnbergi) at Hope Bay, Antarctic Peninsula: differences between breeders and non-breeders. Polar Biol. 2019 Feb;42(2):385–94.

49. Doidge DW, Croxall JP. Factors affecting weaning weight in Antarctic fur seals Arctocephalus gazella at South Georgia. Polar Biol. 1989 Jan 1;9(3):155–60.

50. Nagel R, Mews S, Adam T, Stainfield C, Fox-Clarke C, Toscani C, et al. Movement patterns and activity levels are shaped by the neonatal environment in Antarctic fur seal pups. Sci Rep. 2021 Jul 12;11(1):14323.

51. da Silva J, Terhune JM. Harbour seal grouping as an anti-predator strategy. Anim Behav. 1988 Sep 1;36(5):1309–16.

52. Helfman GS. Threat-sensitive predator avoidance in damselfish-trumpetfish interactions. Behav Ecol Sociobiol. 1989 Jan 1;24(1):47–58.

53. Wood TC, Moore PA. Fine-tuned responses to chemical landscapes: crayfish use predator odors to assess threats based on relative size ratios. Ecosphere. 2020;11(9):e03188.

54. Ferrari MCO, Trowell JJ, Brown GE, Chivers DP. The role of learning in the development of threat-sensitive predator avoidance by fathead minnows. Anim Behav. 2005 Oct 1;70(4):777–84.

55. Palmer MS, Packer C. Reactive anti-predator behavioral strategy shaped by predator characteristics. PLOS ONE. 2021 Aug 18;16(8):e0256147.

56. De Vos A, Justin O’Riain M, Meyer MA, Kotze PGH, Kock AA. Behavior of Cape fur seals (*Arctocephalus pusillus pusillus*) in relation to temporal variation in predation risk by white sharks (*Carcharodon carcharias)* around a seal rookery in False Bay, South Africa. Mar Mammal Sci. 2015 Jul;31(3):1118–31.

57. Suraci JP, Smith JA, Chamaillé-Jammes S, Gaynor KM, Jones M, Luttbeg B, et al. Beyond spatial overlap: harnessing new technologies to resolve the complexities of predator–prey interactions. Oikos. 2022;2022(8):e09004.

58. Paijmans AJ, Berthelsen AL, Nagel R, Christaller F, Kröcker N, Forcada J, et al. Little evidence of inbreeding depression for birth mass, survival and growth in Antarctic fur seal pups. Sci Rep. 2024 Jun 1;14(1):12610.

59. Pollet IL, Arnyek A, Baak JE, Clark R, Comeau-Ouellette J, Grewal AC, et al. Technological advancements: a global review of the use of camera technology in wildlife research. Environ Rev. 2025 Jan;33:1–14.

60. Zampetti A, Mirante D, Palencia P, Santini L. Towards an automated protocol for wildlife density estimation using camera-traps. Methods Ecol Evol. 2024;15(12):2276–88.

61. Chimienti M, Kato A, Seydi V, Schoombie S, Hinke JT, Joy R, et al. Reviewing seas of data: Integrating image-based bio-logging and artificial intelligence to enhance marine conservation. Methods Ecol Evol [Internet]. [cited 2026 Jan 7];n/a(n/a). Available from: https://onlinelibrary.wiley.com/doi/abs/10.1111/2041-210X.70063

