## Supplementary Information for "Fine-scale spatiotemporal predator-prey interactions in an Antarctic fur seal colony"

### Supplementary Information for the manuscript titled “Fine-scale spatiotemporal predator-prey interactions in an Antarctic fur seal colony”

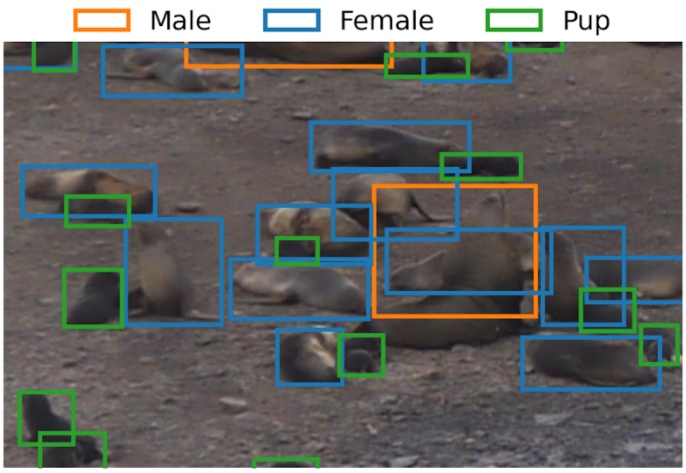


**Supplementary figure 1:** representative example of a crowded beach section showing ground truth bounding boxes. Such dense clusters occur frequently in the field, necessitating the use of overlapping annotations to distinguish individual animals (Male: orange, Female: blue, Pup: green) even when they are partially occluded or resting against one another.

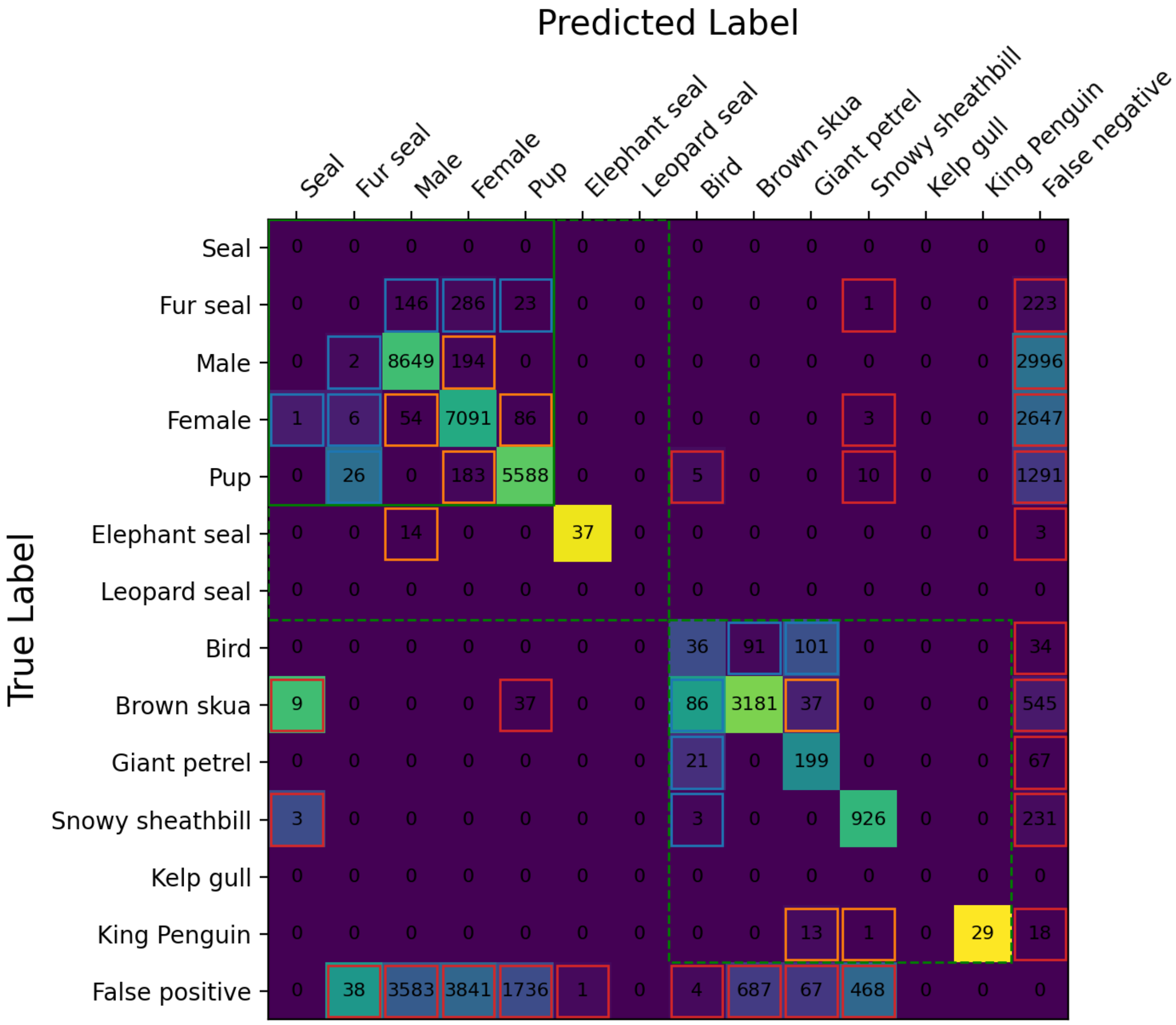


**Supplementary figure 2:** Confusion matrix for the test set across all classes. The vertical axis represents the true labels, while the horizontal axis indicates the labels predicted by the neural network. Objects that were not detected by the network are listed in the 'False negative' column, while the 'False positive' row includes background regions incorrectly classified as objects. Cell numbers denote raw instance counts, and the heatmap intensity reflects precision (column-normalized), where lighter colors (yellow) indicate that a high percentage of predictions for a given class were correct. Colored outlines categorize error types: blue boxes highlight hierarchical discrepancies where a generic label was confused with a specific one (e.g., ground truth '*bird*' vs. predicted 'Brown skua') or vice versa; orange boxes indicate intra-taxa errors between species of the same group (e.g., confusing a brown skua with a giant petrel); and red boxes denote inter-taxa errors across biological orders (e.g., confusing a *seal* with a *bird*). Dashed green lines delineate the main taxa (*seals* vs. *birds*), while the solid green line marks the fur seal subgroup.
We can use this matrix to calculate F1 scores for each class.

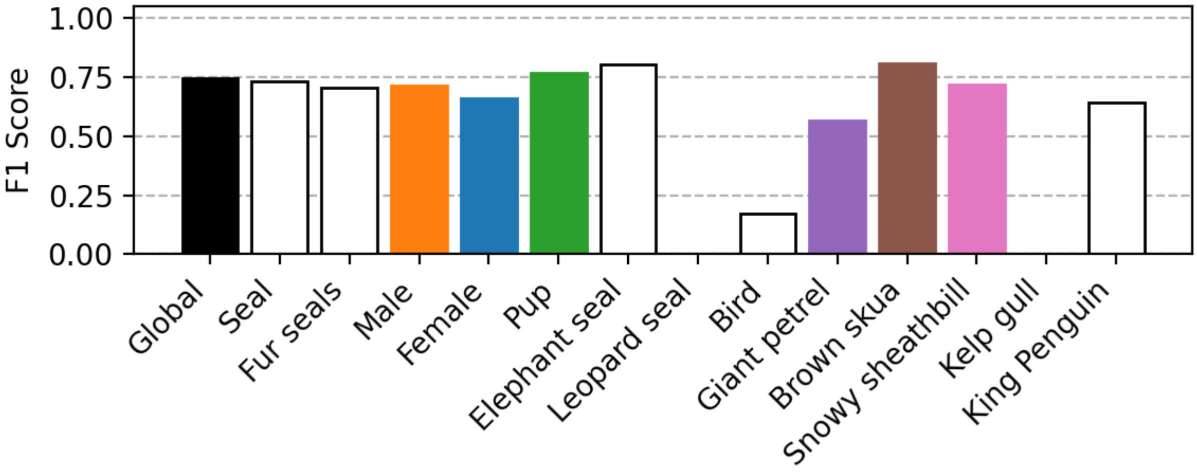


**Supplementary figure 3:** The diagram displays the global F1 score and the F1 score per class, with the six focal classes highlighted in colour. The observed variation in detection performance across these classes is driven primarily by three interacting factors: visual contrast against the background, the number of training annotations (dataset abundance), and object resolution (size relative to the frame). Note that while kelp gull and leopard seal are listed, no scores are reported for them; as infrequent visitors to the study area, these species were not the primary focus of the investigation and consequently had no instances in the test set to evaluate.

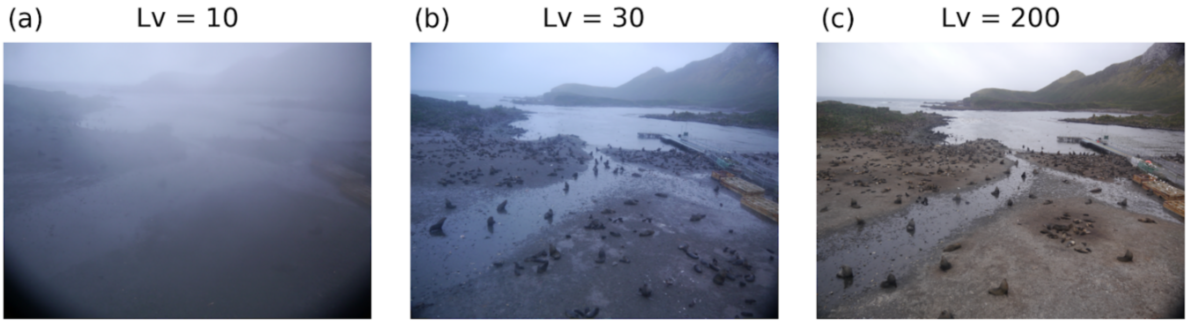


**Supplementary figure 4:** Example images of the Freshwater beach site illustrating varying sight conditions and field of view (FOV). Image clarity is quantified using the Laplacian variance (Lv) metric, where higher values correspond to sharper, clearer images. (a) Low visibility conditions (Lv = 10) caused by environmental factors such as fog or water droplets accumulating on the lens. (b) Moderate visibility (Lv = 30). (c) High visibility with clear atmospheric conditions (Lv = 200). The Lv metric was utilized to remove images with insufficient clarity or severe obstructions from the large number of images in the dataset. By manually assigning classification labels for a subsample of images, we could define a classification threshold for the automated dataset which maximizes the F1 score in this classification problem. Our filtering threshold was set to 30.

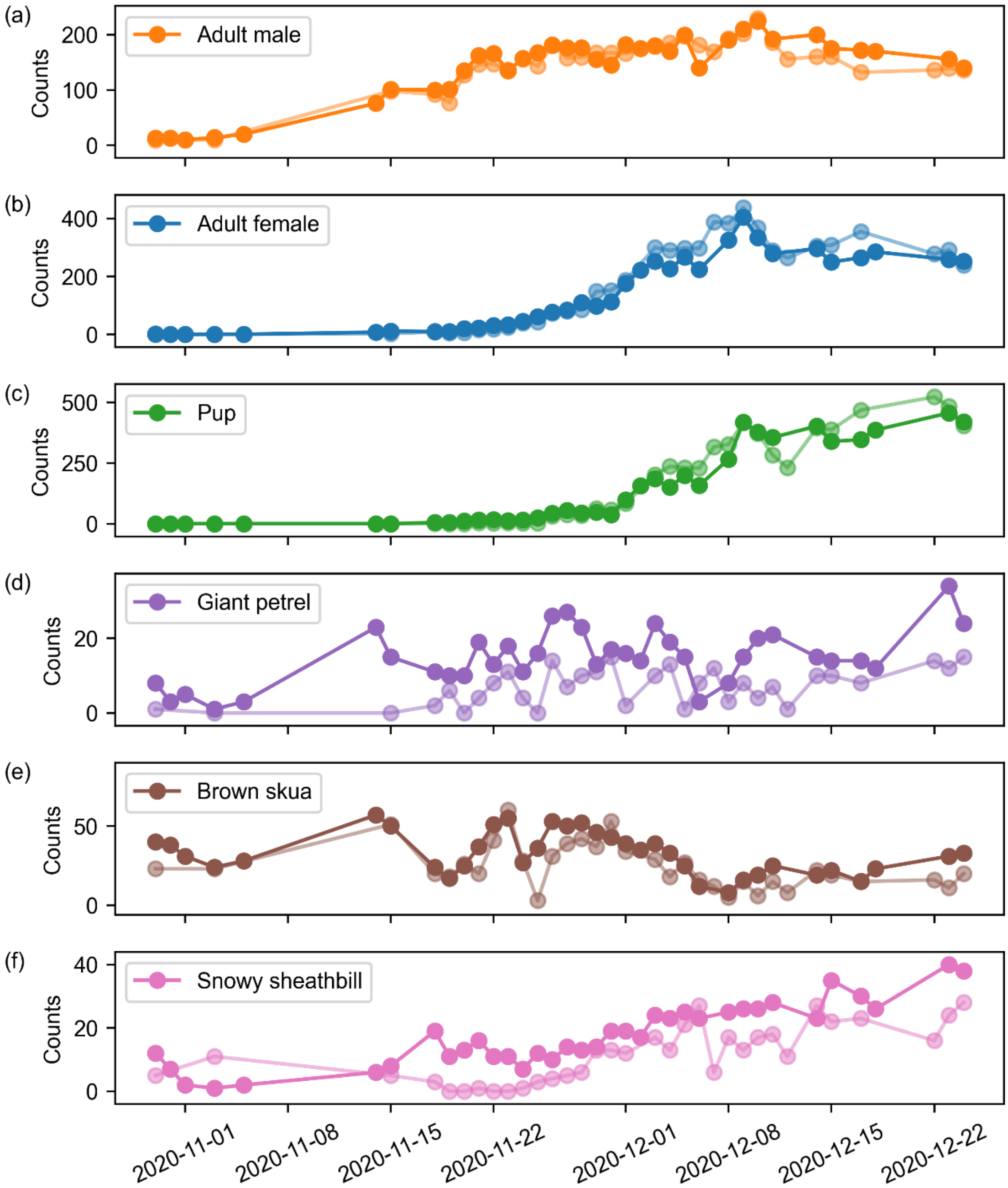


**Supplementary figure 5:** Daily maximum counts from a neural network automatic detection algorithm (solid colours) compared with manual counts (semi-transparent colours) for Antarctic fur seals: (a) adult males, (b) adult females and (c) pups, as well as the predator species: (d) giant petrels, (e) brown skuas and (f) snowy sheathbills.

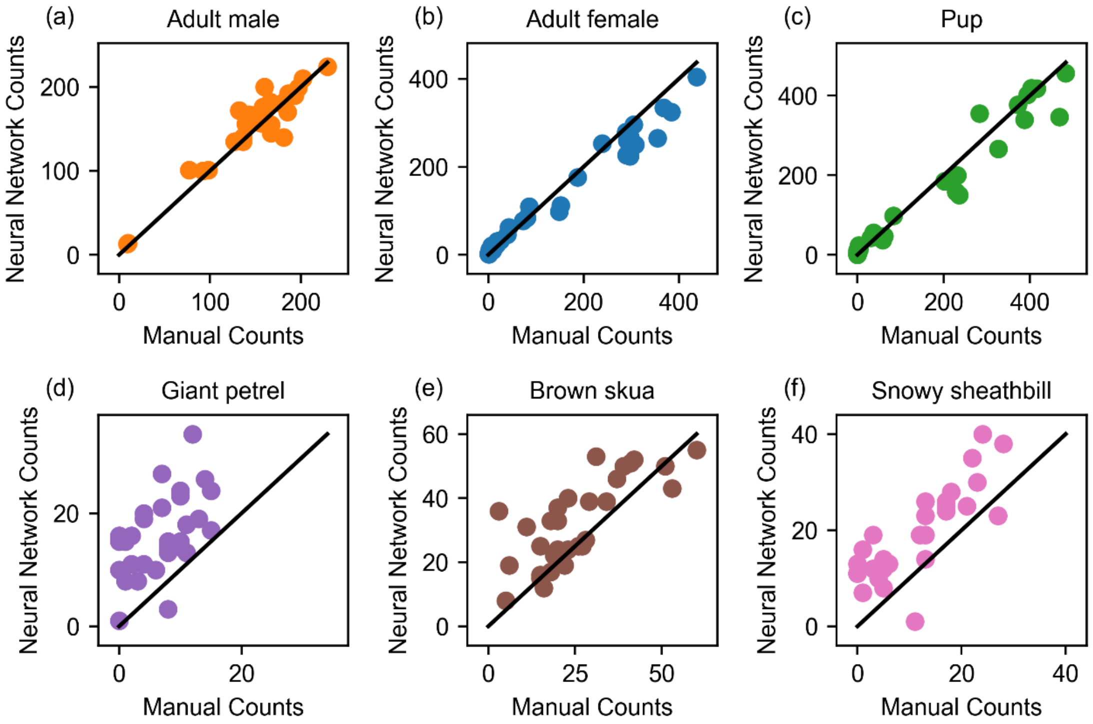


**Supplementary figure 6**: Comparison of the daily maximum neural network counts with the manual counts for each category of Antarctic fur seal: (a) adult male, (b) adult female, and (c) pup, as well as the avian predator species (d) giant petrel, (e) brown skua, and (f) snowy sheathbill. The black lines indicate the reference for a one-to-one agreement between the two counting methods.
